# Characterization of a major QTL for sodium accumulation in tomato shoot

**DOI:** 10.1101/2024.05.21.595121

**Authors:** A. Héreil, M. Guillaume, R. Duboscq, Y. Carretero, E. Pelpoir, F. Bitton, C. Giraud, R. Karlova, C. Testerink, R. Stevens, M. Causse

**Affiliations:** INRAE, UR1052 GAFL, 84143 Montfavet, France; GAUTIER Semences, Route d’Avignon, 13630 Eyragues, France; INRAE, UE A2M, 84143 Montfavet, France; Laboratory of Plant Physiology, Wageningen University, 6700 AA Wageningen, The Netherlands

**Keywords:** Salt stress, GWAS, *Solanum lycopersicum*, HKT1, breeding

## Abstract

Soil salinity is a serious concern for tomato culture, affecting both yield and quality parameters. Although some genes involved in tomato salt tolerance have been identified, their genetic diversity has been rarely studied. In the present study, we assessed salt tolerance-related traits at juvenile and adult stages in a large core collection and identified salt tolerance QTLs by genome-wide association study (GWAS). The results suggested that a major QTL is involved in leaf sodium accumulation at both physiological stages. We were able to identify the underlying candidate gene, a well-known sodium transporter, called SlHKT1.2. We showed that an eQTL for the expression of this gene colocalized with the sodium content QTL. A polymorphism putatively responsible for its variation was identified in the gene promoter. Finally, to extend the applicability of these results, we carried out the same analysis on a test-cross panel composed of the core collection crossed with a distant line. The results indicated that the identified QTL retained its functional impact even in a hybrid genetic context: this paves the way for its use in breeding programs aimed at improving salinity tolerance in tomato cultivars.

**Summary statement:** The genetic diversity of sodium accumulation in tomato shoot under salt stress is associated with an expression QTL of *Sl*HKT1.2. This gene, already identified as involved in tomato salinity tolerance, has been differentially selected during domestication and is of interest for the development of salinity-tolerant varieties.

## 1. Introduction

Soil salinity is a major agricultural challenge: excess soil salinity occurs under all climatic conditions and results from both natural events and human-induced actions. Saline soils occur in arid and semi-arid regions where rainfall is insufficient to meet water requirements of crops and leach mineral salts away from the root zone. Precise world statistics on the global extent of soil salinisation does not exist, but many sources claim that between 25% and 30% of irrigated lands are affected by high salt and are commercially unproductive. Furthermore, salinity is expected to increase in many areas of the world due to global warming [1]. From an economic point of view, the cost of salt-induced soil degradation is difficult to estimate, as it depends on many factors, such as the nature of crops, the extent and severity of soil degradation, the effectiveness of soil and water management practices, but it is estimated at an average cost of $441 ha^−1^ [2].

From a biological perspective, the adverse effects of salinity on plant growth arise primarily because of the increase in osmotic pressure, triggered by excessive Na^+^ accumulation, causing a subsequent decline in the soil’s water potential, inhibiting root water absorption and consequently diminishing water availability. Beyond osmotic stress, an altered cytoplasmic Na^+^/K^+^ balance impairs the functionality of a myriad of K^+^-dependent enzymes [3]. Finally, the cumulative effects of salt-induced osmotic stress and ionic disequilibrium compromise photosynthesis, perturbing the plant’s energy metabolism. These conditions may catalyse the generation of reactive oxygen species (ROS), instigating oxidative degradation within cellular structures [4]. Salinity tolerance in plants is not a ubiquitous trait but varies between and within species. This variability provides fertile ground for the identification of specific genetic loci and natural variations which can play an essential role in improving salt stress tolerance. The first studies, carried out mainly on the model plant *Arabidopsis thaliana*, identified a plethora of genes involved in salt stress response [3]. However, this species exhibits relatively low salt tolerance and is not easily phenotyped for relevant agronomic traits such as yield. In contrast, tomato (*Solanum lycopersicum*) stands out as an agriculturally valuable species, and its role as a model organism for various biological processes, including abiotic stress tolerance, has been gaining momentum. Native to western South America, numerous wild relatives of tomato, such as *S. pennellii*, *S. chilense*, and *S. cheesmaniae*, are adapted to saline environments. Notably, these species have demonstrated higher salinity tolerance than *S. lycopersicum* [5–7], indicating a promising avenue for identifying salt resistance genes and quantitative trait loci (QTLs).

Studies of the genetic architecture underlying salt tolerance in tomato have mainly relied on classical methodologies, notably QTL mapping on biparental populations [8,9,10]. Such approaches inherently limit the discovery of variants present in only one or a limited set of wild donor genotypes. However, by embracing the potential of Genome-Wide Association Studies (GWAS) using a diversity panel encompassing a large number of accessions, a more comprehensive understanding of the genetic landscape can be obtained. Such an approach was successfully implemented to analyse the root Na^+^/K^+^ ratio in a diverse panel of tomato genotypes [11]. In this study, the authors identified and validated a HAK/KUP/KT transporter, named *Sl*HAK20, which plays a pivotal role in the Na^+^/K^+^ ratio in roots. Furthermore, natural variation in the components of the SOS-pathway, specifically *Sl*SOS1 and *Sl*SOS2, has been associated with the root Na^+^/K^+^ ratio in tomato within the same panel [12,13].

In the present study, we introduced a novel GWAS for tomato salt tolerance, with an emphasis on mineral content traits, primarily Na^+^ and K^+^. These elements were analysed in different organs at different physiological stages. We used two distinct but related panels : an inbred core collection and an F1 test-cross panel derived from the former [14]. This second panel aimed to evaluate the applicability and relevance of the QTLs identified at the hybrid level. The first part of this work is dedicated to trait analysis, by elucidating their significance in salt tolerance. Then, a GWAS was carried out, revealing several potential candidate genes. A major QTL hotspot associated with Na^+^ content was identified on chromosome 7, with two underlying candidate genes previously studied, *Sl*HKT1.1 and *Sl*HKT1.2 [15]. Here we showed that this association also co-localised with a root expression QTL for *Sl*HKT1.2. Finally, The analysis of the evolution of the haplotypes of this gene suggested that domestication may have resulted in the fixation of a more highly expressed variant of *Sl*HKT1.2.

## 2. Materials and methods

### 2.1. Plant materials and growth conditions

Two panels of tomato genotypes were used to analyse the genetic response to salt stress. The first panel was composed of 166 homozygous lines (mainly cherry tomato accessions and 15 *S. pimpinellifolium* accessions) from a core collection previously characterised in other studies [16,17]. We will refer to this first panel as CC (Core Collection). The second panel was composed of 144 F1 hybrids obtained by crossing 144 lines of the CC with a large-fruited line Ferum-TMV (FTMV) used as a tester. We will refer to this panel as TC (Test Cross). The list of accessions is provided in **Supplementary Table 1**.

### 2.2. Growth conditions and measurement of biomass/agronomic traits

The CC panel has been grown in two experiments, one in hydroponic condition in a phytotron where plants were grown for 20 days, another in greenhouse under agronomic conditions. The TC panel has been studied only in greenhouse condition.

#### Phytotron trial

Temperatures were set at 24°C (day) /18°C (night), with a photoperiod of 16/8h and 65% relative humidity. Light was supplied by cool white fluorescent lamps at a photosynthetic photon flux density of 165 μmol m^−2^ s^−1^ at the leaf level. The hydroponic setup consisted of opaque plastic containers (50 L) covered by a drilled opaque lid with 20 holes (d= 1.43 cm) at regular intervals. A 2-cm-long rockwool plug holder was placed in each hole. Containers were filled with a standard nutrient solution (Liquoplant Rose, Plantin, Courthézon, France) diluted to 4% (2.0 mS) and adjusted to pH 5.7 using hydrochloric acid. The solution was renewed every 5 days. The seeds were sown directly in rockwool plug holders and germinated on a heated tray (25°C). At 7 days after germination (d.a.g), seedlings were transferred to the hydroponic system. The salt-stress condition was initiated at 10 d.a.g by adding 60 mM NaCl to the nutrient solution. The salinity was then increased to 120 mM at 12 d.a.g. Another group of plants was used as controls (without any addition of NaCl). The design of the experiment was structured as follows: It comprised 24 containers, each holding 32 plants. Among these containers, 12 were used for control condition, while the other 12 were used for stress condition. The plants were arranged in a random but symmetrical layout, which facilitated simultaneous sampling of the two sets of plants and enabled plasticity traits to be calculated by minimising spatio-temporal effects. Also, two replicates *per* genotype were placed side by side in each container. The experiment was carried out twice to obtain between 4 and 6 biological replicates per genotype for each condition. Plant harvesting started at 20 d.a.g and was carried out over a 3 h time window. For each replicated genotype grown in the same container, whole roots from two plants were weighted, washed with deionised water, pooled, snap-frozen in liquid nitrogen, and stored at −80°C for later analysis. The shoot parts were also weighted, pooled, washed with deionised water and dried in a forced-air oven (70°C) for 48h, leading to fresh weight (FW) and dry matter content. (DMC).

#### CC greenhouse trial

The CC collection was grown from March to July 2021 in a glasshouse at INRAE GAFL, Avignon, under soilless conditions in rockwool. Plants were irrigated with a standard fertigation solution (Liquoplant Rose, Plantin, Courthézon, France) where NaCl was added, resulting in a final EC of approximately 8 dS.m-1. In order to gradually adapt plants to salt stress, the introduction of NaCl into the fertigation solution began one week after the plantation. Then, 50 mmol L^−1^ of NaCl was added for another week before increasing to the final level (80 mM). For each accession of the CC collection, the experiment was conducted with three biological replicates. Plant height (in cm), stem diameter (in mm) and leaf length (in cm) under the 4th truss were measured. Red-ripe fruits were harvested twice a week to assess mean fruit weight.

#### TC greenhouse trial

The TC panel was grown in a greenhouse at Gautier Semences, Eyragues, France, twice in 2021, first in spring under control condition, then in autumn under salt stress, using the same protocol as for the CC, but in addition fruit yield was assessed.

### 2.3. Mineral content measurement

Two fully developed leaves were harvested on each plant above the fifth flower cluster prior to the ripening of the first cluster. Petioles were stripped of their leaf blades and promptly stored at −20°C until analysis. Petiole sap was extracted using an extraction bag and a semi-automated homogenizer (HOMEX 6; Bioreba, Reinach, Switzerland). The filtered petiole sap obtained was utilized to determine the sodium (Na^+^) and potassium (K^+^) content using an ion-selective electrode meter specific for each ion following the manufacturer’s instructions (LAQUAtwin Na-11 and K-11; Horiba Instruments Inc., Irvine, CA). For the evaluation of Na^+^ and K^+^ levels in fruits, a minimum of five red ripe fruits were collected from the third to the sixth truss and pooled for each replication. Subsequently, Na^+^ and K^+^ content was assessed on crushed fruit pericarps using the same device. For ionome measurement in young plants, dried leaf samples were ground, weighted, and digested with nitric acid and hydrogen peroxide using a block digester. Analysis of Na, K, Mg, Mn, Ca, Fe, and Cu levels was conducted via MP-AES (Microwave Plasma Atomic Emission Spectrometry) (Agilent 4200MP-AES, Santa Clara, CA) at the BPMP laboratory (Montpellier, France).

### 2.4. Estimation of genotype means and heritabilities

Prior to association studies, phenotypic data were aggregated: for phenotypes whose repetitions were carried out at different times (i.e., all fruit derived phenotypes: Na^+^ and K^+^ fruit content), best linear unbiased predictors (BLUPs) were estimated using the R lme4 package to correct for the harvest-day effect:

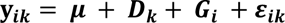

Where ***μ*** is the overall mean, ***G_i_*** represents the i^th^ genotype effect, ***D_k_*** represents the k^th^ harvest-day effect and ***ε_ik_*** the ik^th^ residual variation with 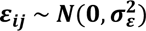.

Biomass traits from the greenhouse trial (measured on a single day) were aggregated by a simple mean. Biomass and leaf mineral contents measured in the phytotron were also corrected using a mixed effect model:

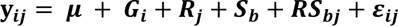

Where ***R_j_*** represents the j^th^ replicate effect, **S*_b_*** is the b^th^ block effect, ***RS_bj_*** is the bj^th^ block × replicate interaction effect with 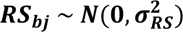.

For traits measured on seedlings under control and salt stress conditions, a random-effect analysis of variance was conducted on the whole population to test for genotype (**G**), treatment (**T**), and their interaction (**G×T**) effects with the following model using R/lme4:

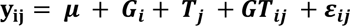

Where **T_j_** is the effect of the **j^th^**treatment, and ***GT_bj_*** is the ij^th^ genotype × treatment interaction effect with 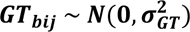. Broad-sense heritability was calculated from the above model according to [18]:

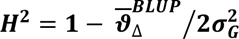

Where 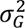 is the genotype variance, 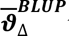 the average standard error of the genotypic BLUPs.

Phenotypic plasticity traits were computed as the ratio of stress to control values. This index was then used as a phenotype *per se* for QTL analysis. We will thus refer to (C), (S) and (D) for traits measured in control, stress and plasticity conditions. When all the genotypes were taken into account, the average effect of the stress was reported as the mean relative difference between control and stress conditions and converted into the percentage increase or decrease due to the stress. The significance of the treatment effect was then calculated by using a likelihood ratio test from R/lmtest, by comparing the goodness-of-fit between two models, one considering the treatment effect and a second one not taking it into account. Prior to GWAS analyses, we applied a Box-Cox transformation of the means calculated from replicates to correct for heteroscedasticity and non-normality of error terms [19]. Trait means of all accessions and treatments are presented in **Supplementary Tables 2 and 3.**

### 2.5. Genotypic data and GWAS analysis

To detect associations between genotypic values and genomic markers, GWAS was performed using a set of 6,251,927 SNPs from genome resequencing. All procedures used for alignment, SNP calling and post processing are described in [20]. SNP positions refer to the Heinz_1706 v.4.0.0 reference genome. A univariate GWAS was performed by implementing the following linear mixed model in R/GENESIS [21]:

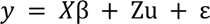

where *y* is the vector of phenotypic means, *X* is the molecular marker score matrix, β is the vector of marker effects, Z is an incidence matrix, *u* is the vector of random background polygenic effects with variance σ^2^_u_ = K σ^2^_G_ (where K is the kinship matrix, and σ^2^_G_ is the genetic variance), and *ε* is the vector of residuals. The null model was fitted using the NullModel() function using only the fixed-effect covariates. We included the first three eigenvectors estimated from the PCA-based overall genotypic matrix using PLINK 2.0 [22]. Single-variant association tests were performed with the assocTestSingle() function using the Average Information REML (AIREML) procedure to estimate the variance components and score statistics. We then used an arbitrary genome-wide significance thresholds of LOD = 6 for SNP significance. To estimate the proportion of variance explained (PVE) by lead SNP, we used the following formula proposed by [23] using outputs from R/GENESIS:

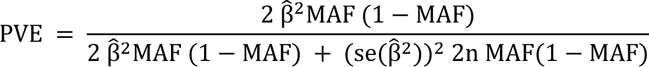

Where 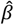 is the effect size, 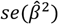 is the standard error for the effect size, *MAF* the minor allele frequency and *n* the sample size.

### 2.6. Haplotype and diversity analysis

Due to the low number of *S. pimpinellifolium* accessions in the CC panel, we merged this panel with the Varitome core collection [24] to have a broader view of tomato genetic diversity. This led to a panel of 331 resequenced accessions. Using this combined panel, we performed different haplotype analyses. A haplotype heatmap was generated using the R/pheatmap, using Ward distance for clustering the genotypes (rows). Cluster number for *k*-means was calculated according to the elbow method using a screen plot with the identification of the optimal number of clusters when the total intra-cluster variation was minimized. The multiple mean comparison to test significant differences among haplotype clusters was conducted using a linear model. We used R/emmeans to calculate the p-value of pairwise comparisons among clusters and R/multcompView to display the Tukey test, fixing the significance threshold at 0.05. To investigate if *Sl*HKT1.2 was under selection we computed allele-specific extended haplotype homozygosity (EHH). This analysis was conducted both upstream and downstream of the most significant SNP in *Sl*HKT1.2 promotor sequence using R/rehh. For visualisation, EHH values were integrated across their respective genomic positions to yield an ‘integrated EHH’ (iHH) value for each allele [25].

To identify the conserved regions of *Sl*HKT1.2 promotor within the tomato clade, we performed a sequence alignment involving ten *de novo* assembled genomes of wild tomato relatives *- S. galapagense*, *S. pimpinellifolium*, *S. chmielewskii*, *S. neorickii*, *S. corneliomulleri*, *S. peruvianum*, *S. chilense*, *S. pennelli*, *S. habrochaites*, and *S. lycopersicoides*. These genome sequences were obtained through Hi-C sequencing [26]. For the alignment, the promoter sequence of the *Sl*HKT1.2 gene was extracted from the SL4.0 chromosome 7 sequence between positions 5,126,736 and 5,143,883. Then, the extracted sequence was subjected to BLAST analysis against the ten wild tomato genomes using NOBLAST [27]. Finally, the sequences corresponding to these homologous regions were aligned with the reference SL4.0 sequence using the Needleman-Wunsch pairwise alignment tool, EMBOSS Needle [28].

### 2.7. RNA extraction and RT-qPCR

A total of 92 accessions out of 166 were selected for expression analysis (**Supplementary Table 4**). For each genotype grown in hydroponic condition, whole roots from two seedlings (0.8 to 16 g) grown in the same container were weighted, washed with deionised water, pooled, snap-frozen in liquid nitrogen, and stored at −80°C for later analysis. RNA extraction was performed using the “RNeasy Plant Mini Kit” (Qiagen Sciences, Germantown, MD, USA) following the manufacturer’s protocol, with “On-column DNAse I Digestion Set (Sigma-Aldrich, Saint-Quentin Fallavier, France) to remove any remaining genomic DNA. The amount of RNA was quantified using a Qubit Fluorometer 3.0 and the “Qubit RNA Broad Range Assay Kit” (Invitrogen, Carlsbad, CA, USA). RNA purity (260/230 nm and 260/230 nm ratios) was assessed using the NanoDrop 1000 Spectrophotometer (Thermo Fisher, Waltham, MA, USA); all 260/280 nm ratios were comprised between 1.8 and 2.2 and all 260/230 nm ratios were beyond 2.0. Sample integrity was verified on a Bioanalyzer 2100 (Agilent Technologies, Les Ulis, France) with the “RNA 6000 Nano Kit”; all RNA integrity numbers (RIN) were beyond 7. RNA was then reverse transcribed with the “GoScript Reverse Transcription Mix, Oligo(dT)” (Promega, Madison, WI, USA) and qRT–PCR experiments were performed with the “Brilliant III Ultra-Fast SYBR Green QPCR Master Mix” (Agilent Technologies) according to the manufacturer’s specifications. The primers used for the *Sl*HKT1.1 (Solyc07g014690) were 5′-TCTAGCCCAAGAAACTCAAAT-3′ and 5′-CTAATGTTACAACTCCAAGGAATT-3′, and those for *Sl*HKT1.2 (Solyc07g014680) were 5′-TGAGCTAGGGAATGTAATAAACG-3′ and 5′-AGAGAGAAACTAACGATGAACC-3′, according to [15]. The genes Solyc06g005360 (5′-AAGGGCTTTCCGCTTCATAGT-3′ and 5′-AACTTTCAGCTGGCTCACCAA-3′) and Solyc10g006100 (5′-CCAACTAAAGCGCTGCCACAAC-3′ and 5′-TGGTCCTTGTGTGCTTACTGGC-3′) were used as reference genes. Relative quantification data analysis was determined with the ΔΔCt method.

## 3. Results

### 3.1. Impact of salt stress at juvenile and adult stages

Mean, coefficient of variation as well as percent reduction in mean due to salt stress and variance decomposition of the studied traits are shown in **Table 1**. The impact of salt stress was only characterised for traits measured at juvenile stage (20 dag) as, in greenhouse, the CC panel was grown only in stress condition. We observed a large variation in salt response across the accessions grown in the phytotron: The reduction in total biomass due to salt stress ranged from 13% to 86%. Notably, the disparity in biomass was more pronounced in the shoot, with a decrease of 61.1%, compared to the root, which was reduced by 24.3%, on average. This difference resulted in an elevated shoot-to-root ratio between the two treatments. The variance decomposition of biomass traits remained relatively consistent, with the exception of root biomass, which exhibited an increased residual variance of 40% (**Table 1**). Mineral compositions, however, displayed diverse patterns of variation under salinity stress. Specifically, macro-element concentrations, including calcium (−36%), potassium (−48%), magnesium (−28%), and manganese (−26%), tended to decrease. In contrast, the concentrations of microelements, notably copper and iron, increased by 23% and 74%, respectively. Zinc was the only mineral with no significant variation in concentration under saline conditions. As expected, the sodium content difference between the two treatments was substantial, recording an increase of 2004%. The observed variance in sodium was thus predominantly governed by the treatment effect, accounting for 90.5% of the total variance. Heritability estimates spanned from moderate to high values: for the control conditions, they ranged between 0.55 to 0.73, while under stress conditions, they fluctuated between 0.49 and 0.87. The potassium and sodium contents under saline conditions showed high heritabilities of 0.79 and 0.87, respectively.

**Table 1.**
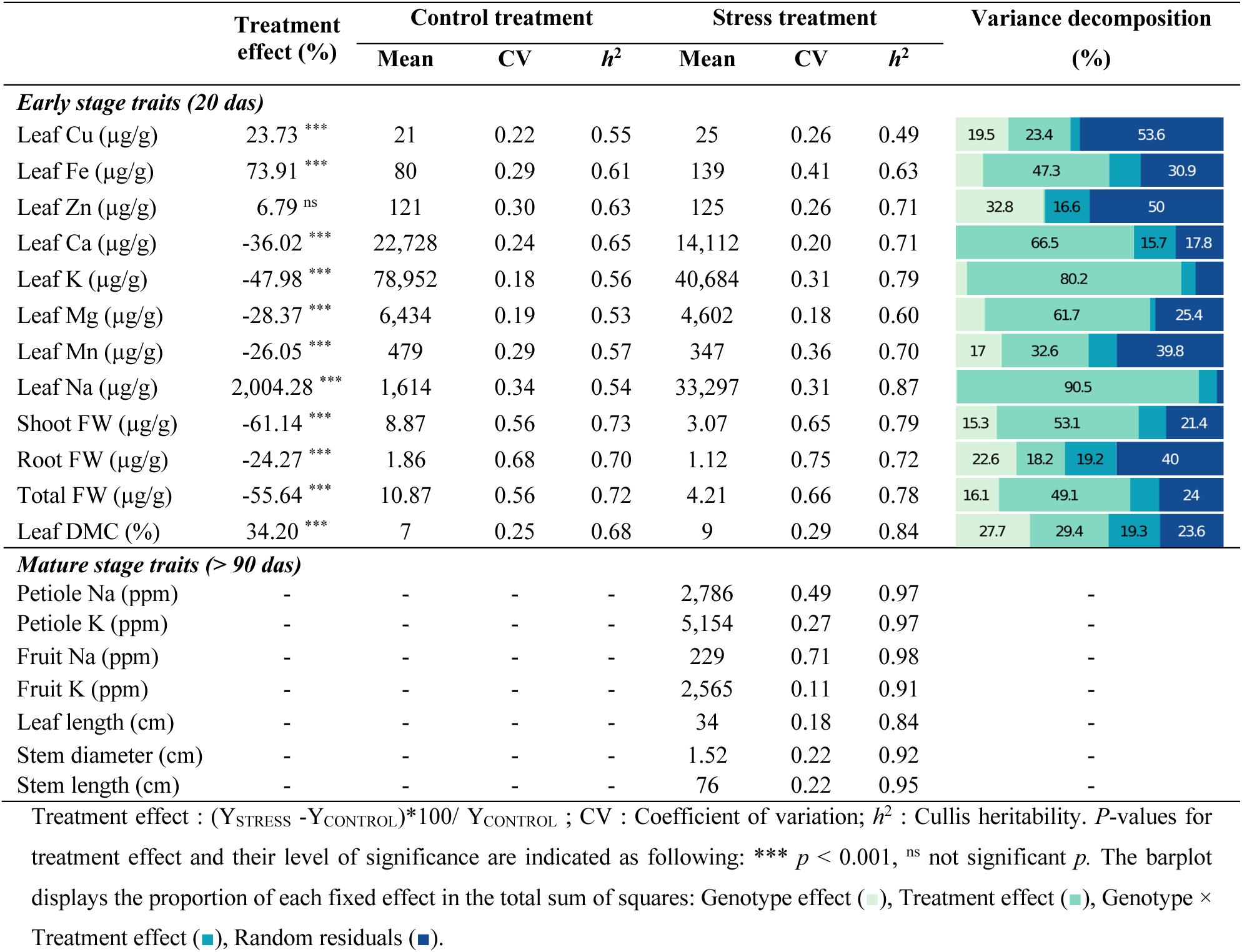
Descriptive statistics and variance decomposition of various physiological and mineral content traits.

The traits evaluated in greenhouse were only characterised under salt stress condition. Broad-sense heritability estimates systematically exceeded those observed during the juvenile stage. This increased heritability could be attributed to two main factors: firstly, the pronounced phenotypic diversity, highlighted by the coefficient of variation reaching 71% (fruit sodium content), and secondly, the experimental design (described in the ‘Materials and Methods’ section) which effectively alleviated potential spatial effects between genotypes.

### 3.2. Correlations of mineral contents with salt tolerance at different stages and in different organs

Based on the correlation and cluster analysis shown in **Figure 1**, distinct groups of correlated traits were identified. All traits relative to Na^+^ content were clustered within the same group (cluster 3), suggesting a common genetic basis for sodium homoeostasis in different organs and at different growth stages. Some traits, such as petiole Na^+^ content, were highly correlated with fruit Na^+^ content (r = 0.78) under stress. It is noteworthy that even with trace concentrations of Na^+^ content in the control condition, there was a substantial correlation between the Na^+^ concentrations of the two treatments (r = 0.63). The traits of cluster 3 were all negatively correlated with the traits in cluster 1, which grouped cation contents (Ca^2+^, Mg^2+^, K^+^) under stress condition and plasticity. Interestingly, the levels of these minerals measured under control conditions (cluster 4) were not correlated with their counterparts under salt stress. Finally, biomass-related traits measured under stress conditions (cluster 1) were correlated only when measured at the same physiological stage. At the adult stage, only the stem diameter exhibited a significant negative correlation with salt content, leaf and stem length being not correlated with Na^+^ content. Stem diameter was also highly correlated with petiole potassium content.

**Figure 1.**
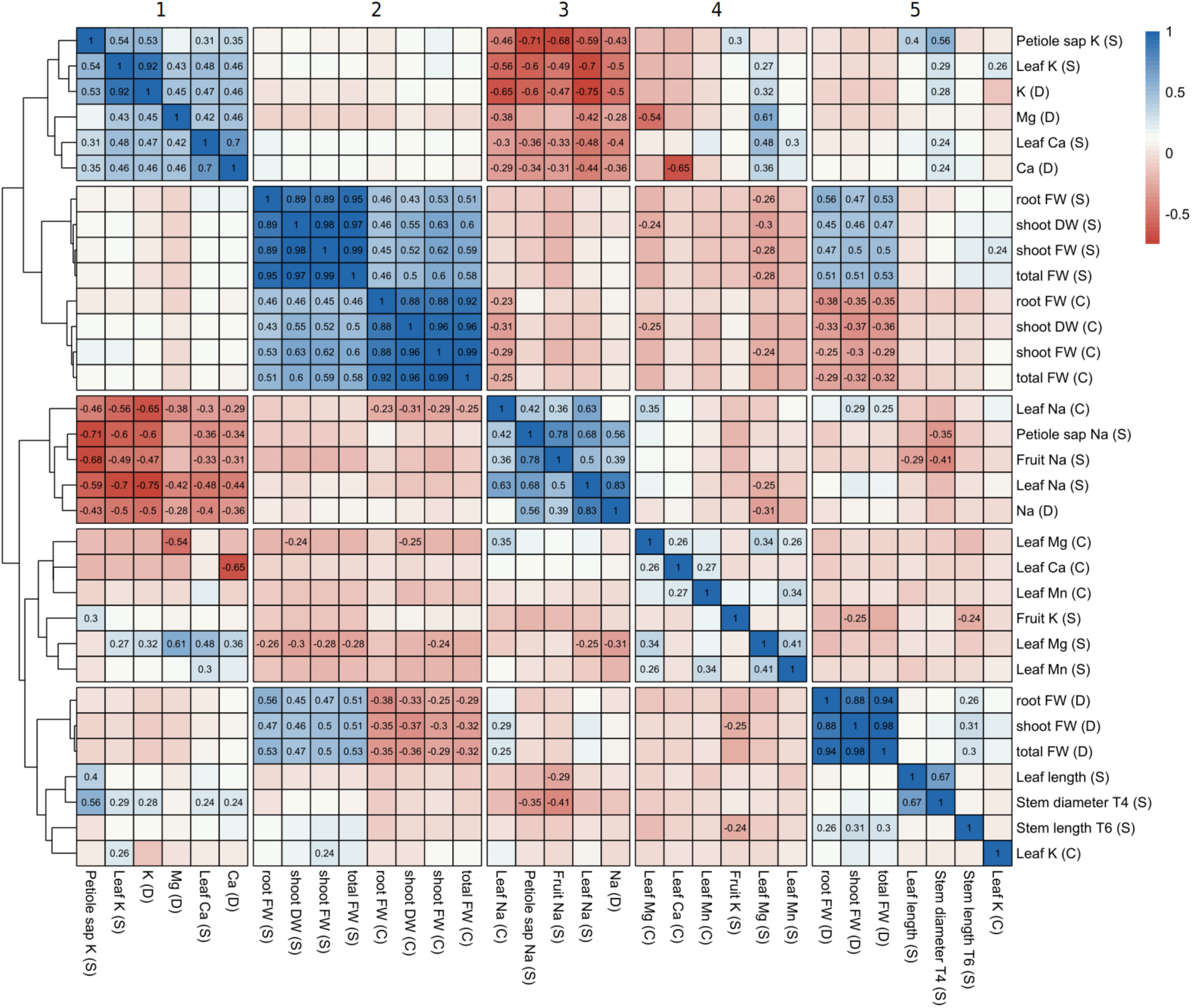
**Correlation between minerals and biomass-related traits.** Hierarchical clustering heatmap of Pearson correlation coefficients between minerals and biomass-related traits. The matrix represents the recorded traits under three conditions: Control (C), Salinity Treatment (S), and the ratio of the two treatments (D). The colour gradient provides insight into the correlation type; a value of 1 means a complete positive correlation (represented by dark red), while a value of −1 indicates a full negative correlation (represented by dark blue) between two traits. Only those correlations that are statistically significant (p-value < 0.01) are shown. Numbers on top of the matrix indicated the clusters of traits.

### 3.3. GWAS analysis reveals a QTL hotspot on chromosome 7

In order to further understand the genetic basis underlying the observed difference in salt tolerance and sodium accumulation, we performed a single-trait GWAS on 166 small-fruited accessions for 22 salt tolerance-related traits (selected based on heritability values and variance decomposition; **Figure 2.a**). We found a total of 1032 SNP that were above the LOD-threshold of 6. Neighbouring SNPs were mostly in linkage disequilibrium. Thus, for subsequent analysis we clustered all markers in linkage disequilibrium, reducing the number of independent QTLs to 14 clusters (**Supplementary Table 5**). A strong association signal for Na^+^ content in leaf (at young stage), fruit and petiole (at adult stage) was identified on chromosome 7, with the lowest p-value of all the traits (LOD >10). In particular, the marker found to be significant for most of the traits was SSL4.0CH07_4555305 (**Figure 2.b**). In this region, more than 7000 SNP markers were significantly associated with at least 11 traits, in the interval ranging from 4,415,544 to 5,127,764 base pairs (bp). This association was found exclusively for sodium levels in each organ type (seedling leaf, mature petiole sap, fruit pericarp), under control and salt stress conditions. This QTL was also found for potassium levels, but exclusively under salt stress conditions. For the traits leaf Na^+^ (S) and petiole Na^+^ (S), this region covered 50 genes, including two High-affinity Potassium Transporters (HKT) genes (*Sl*HKT1.2 (Solyc07g014680) and *Sl*HKT1.1 (Solyc07g014690)), which were both previously identified as playing a pivotal role in sodium homoeostasis in several species, including tomato [15,29]. This QTL also displayed long-range linkage disequilibrium across the chromosome, suggesting that population admixture, epistatic selection, or other evolutionary forces were at work [30]. Beyond this major QTL, few other associations were detected on other chromosomes (**Supplementary Table 5**). Leaf sodium content in stress condition revealed the largest number of associations, with 11 regions identified on 6 chromosomes. We detail below only the associations detected for at least two traits. A significant association was found for leaf sodium content (S), leaf sodium content (D) and leaf Na/K (D) on chromosome 4 (SSL4.0CH04_63269805). The twelve significant SNP for this association were found in one gene, Solyc04g015850, of unknown function but harbouring transmembrane protein characteristics. In this region, at 93 kb, Solyc04g015990 was an interesting candidate gene expressed in roots (**Supplementary Figure 1**). This gene encodes a Cation/H(+) antiporter found to be down regulated by salt stress (fold change in roots: −13.43) [31]. This gene has also been suggested as a candidate for a QTL hotspot for multiple cation xylem content in a RIL population derived from a cross between *S. lycopersicum* and *S. pimpinellifolium* [32]. Another genetic marker (SL4.0CH03_53341434), identified on chromosome 3, exhibited a significant association with four traits related to Na^+^ and K^+^ concentrations at the seedling stage. The most plausible candidate gene responsible for these associations was Solyc03g096670, which encodes a Type 2C protein phosphatase (PP2C). The functional relevance of PP2C in salt tolerance is well documented: especially, its overexpression improved salt tolerance by regulating shoot Na^+^ extrusion [33] or by regulating oxidative stress-related genes [34]. Also, RNA-seq studies revealed that Solyc03g096670 was significantly upregulated under salt stress conditions, both at the root level (7-fold change) [35] and in the leaves (4.8-fold change) [36]. These expression patterns strongly suggest that Solyc03g096670 is involved in the plant adaptive response to salt stress. From the perspective of genetic diversity, substantial variations in gene expression at the root level have been reported. The dataset from Alonge et al., [37] revealed up to a 22-fold change in expression levels between extreme genotypes. This study focused on the impact of structural variants on gene expression and identified a significant association between a deletion located 5 kb downstream of Solyc03g096670 and its expression levels. Such an association suggests the presence of a cis-expression QTL. This hypothesis is further supported by the detection of a Solyc03g096670 eQTL in a panel of introgression lines (ILs) derived from a cross between *S. lycopersicum* ‘M82’ and *S. pennellii*, reported by Ranjan et al., [38]. All these elements suggest that Solyc03g096670 could also be an expression QTL that deserves further analysis. We did not detect any candidate gene for plant weight plasticity at the given threshold (LOD > 6). However, the most significant peak on chromosome 8 (LOD = 5.58) was located at the level of a Late embryogenesis abundant 3 family protein (Solyc08g074720).

**Figure 2.**
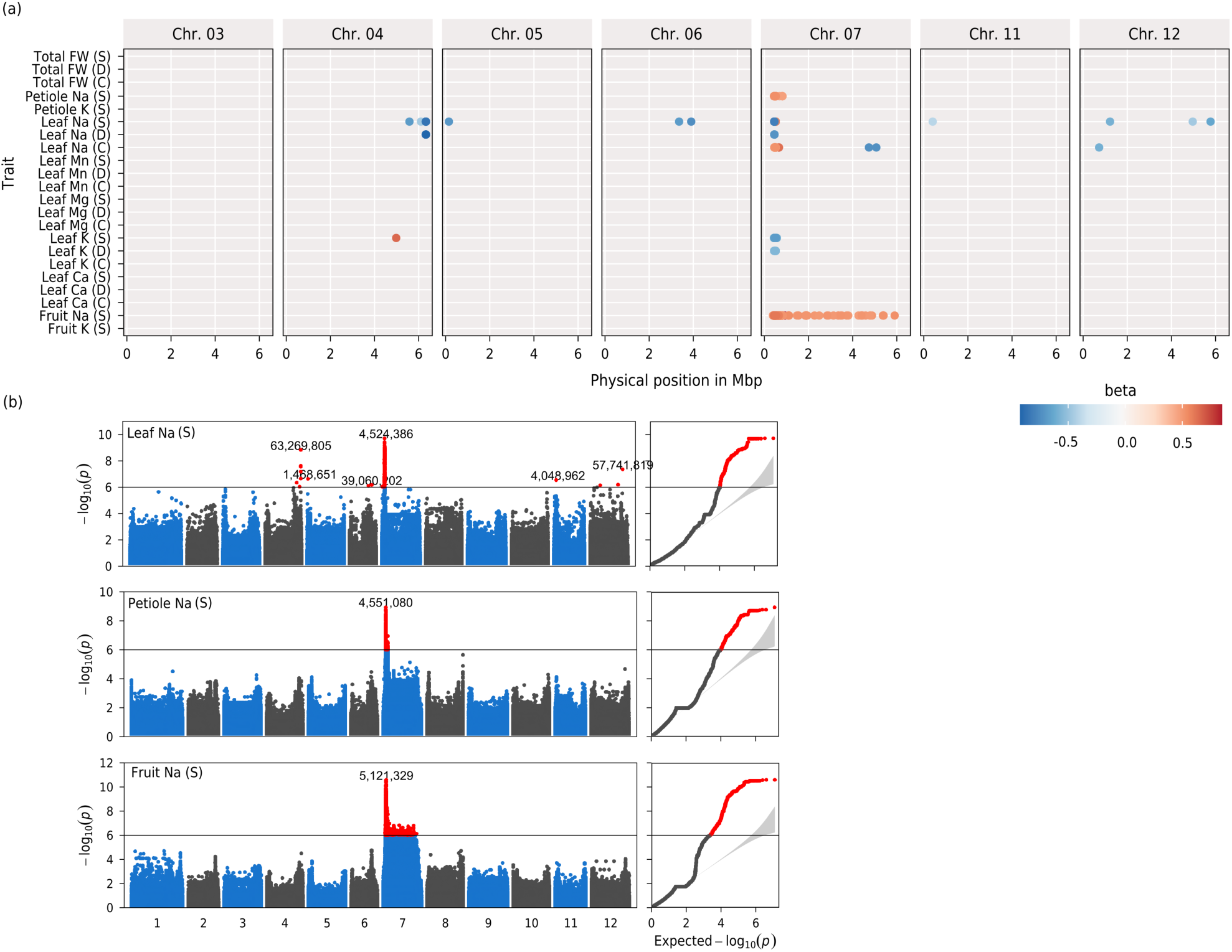
**Chromosome distribution of QTLs associated with biomass and mineral content traits.** (**a**) QTL summary showing the physical positions of GWA significant SNP (LOD > 6). Each row represents the QTL mapping of a single trait. The dot colour denotes regression beta coefficients (blue: negative effect; red: positive effect). The x-axis indicates the physical positions on the SL4.0 tomato genome. In trait nomenclature, C/S/D stands for control, salt and delta conditions (i.e. trait plasticity). (**b**) Manhattan plot and qq-plots for Leaf Na^+^ (S), Petiole sap Na^+^ (S) and Fruit Na^+^ (S). Red dots indicate SNPs that have reached GWAS significance. For ease of interpretation, the position of only the SNP with the highest significant p-value is indicated, along with their respective physical positions.

### 3.4. Candidate gene mining: a focus on HKT genes

The leading SNPs within the QTL on chromosome 7 were close to two HKT1 genes (*Sl*HKT1.1 and *Sl*HKT1.2). Both have been previously identified as playing a pivotal role in sodium homoeostasis in several species, including tomato [15,29]. A first polymorphism analysis revealed that neither of the two HKT1 genes exhibited polymorphisms in the coding sequence that could account for the QTL on chromosome 7. This led us to propose that the underlying polymorphism might be a differential expression of one of these genes. To validate this hypothesis, we measured the expression levels of both HKT1 genes in root samples exposed to salt stress in phytotron, given that both genes are predominantly expressed in roots (**Supplementary Figure 1**). Our analysis revealed a substantial difference in *Sl*HKT1.2 expression between accessions harbouring the reference and alternative alleles. A major association was identified for *Sl*HKT1.2, colocalizing with the previously mentioned QTL on chromosome 7 (**Figure 3.a**). This QTL accounted for a substantial portion of the variance in *Sl*HKT1.2 gene expression, explaining approximately 50% of the variance of this gene expression (**Figure 3.b**; t-test p-value = 3.6 10^−11^). Conversely, no significant differences were detected in *Sl*HKT1.1 expression. Furthermore, *Sl*HKT1.2 expression was negatively correlated with leaf sodium content (r = −0.63, **Figure 3.d**) and positively correlated with potassium content (**Figure 3.e**; r = 0.5). No significant correlation was observed for *Sl*HKT1.

**Figure 3.**
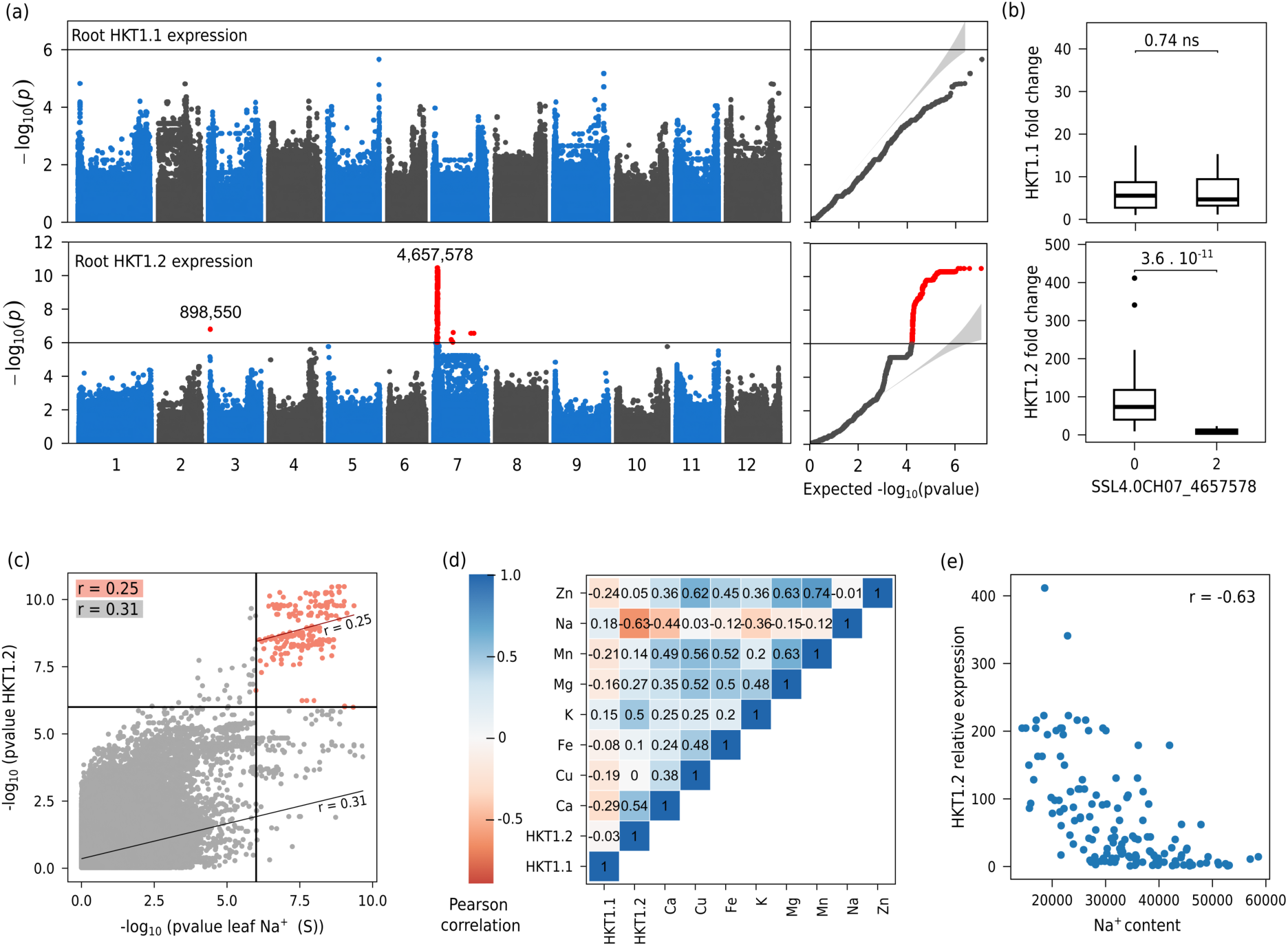
**Colocalisation between sodium content QTL and expression QTL (eQTL) for *Sl*HKT1.2 root expression** (**a**) Manhattan plot and qqplot of GWAS analysis of *Sl*HKT1.1 and *Sl*HKT1.2 expression in root tissues under salt stress. (**b**) Boxplot displaying the difference in *Sl*HKT1.1 and *Sl*HKT1.2 expression between the three genotypes of SSL4.0CH07_4657578 (0 reference alleles, 1 heterozygous, 2 alternative alleles). (**c**) Scatterplot displaying the correlation between GWAS p-values for the Leaf Na^+^ (S) and GWAS p-values for *Sl*HKT1.2. Significant markers for both traits are colored in red. (**d**) Correlation matrix displaying the correlations between the expression of the two HKT1 genes in root tissues with leaf mineral content. (**e**) Relationship between leaf sodium content and *Sl*HKT1.2 relative expression.

As we found a colocalisation between the sodium content QTL and the eQTL for *Sl*HKT1.2 expression, we searched for polymorphisms that could explain this difference in gene expression. We thus first carried out a local reanalysis of *Sl*HKT1.2 expression association, using in addition to SNP, indels and short structural variants detected by the ANNOVAR algorithm (**Figure 4**). In the promoter region, we detected few significant polymorphisms, including three SNPs and one short indel (T/TC) at 1,564 bp from the initiation start. We also compared the Heinz1706 SL4.0 reference genome to the one of BGV007931, a *de novo* genome assembly of a Mexican *S.l. cerasiforme* tomato with a level of *Sl*HKT1.2 expression close to zero, compared to the reference genome accession Heinz1706 [37]. The comparison of both sequences revealed additional interesting polymorphisms, including two short tandem repeats at 2,491 and 2,680 bp from the initiation site and of lengths 10 and 43 bp, respectively (**Figure 4**). As these two polymorphisms could not be captured with our short-read sequencing, we did not include them in the GWAS analysis. Finally, we sought to elucidate the potential impact of the mutations that target the HKT1.2 promoter sequence, focusing on the alteration of promoter motifs. We only considered polymorphisms that were significant in GWAS. We identified four candidate polymorphisms located in the promoter region. Among them, two motifs were predicted to be structurally affected by mutations: a nucleotide substitution at position −972 of the transcription start site (TSS) and an insertion at position −1,564 bp (**Table 2**). Both markers were in complete linkage disequilibrium (r^2^ = 1). The insertion at position −1,564 bp is particularly interesting, as the alternative allele disrupts an ABRE (ABscisic acid-Responsive Element) motif. ABRE motifs have already been identified in many HKT1 gene promoters in several species, including *Arabidopsis*, wheat and rice [39]. Furthermore, in *Arabidopsis*, a single ABA-responsive element in the *At*HKT1;1 gene promoter has been shown to be essential for gene expression, particularly in response to sodium chloride (NaCl) and abscisic acid (ABA) [40].

**Figure 4.**
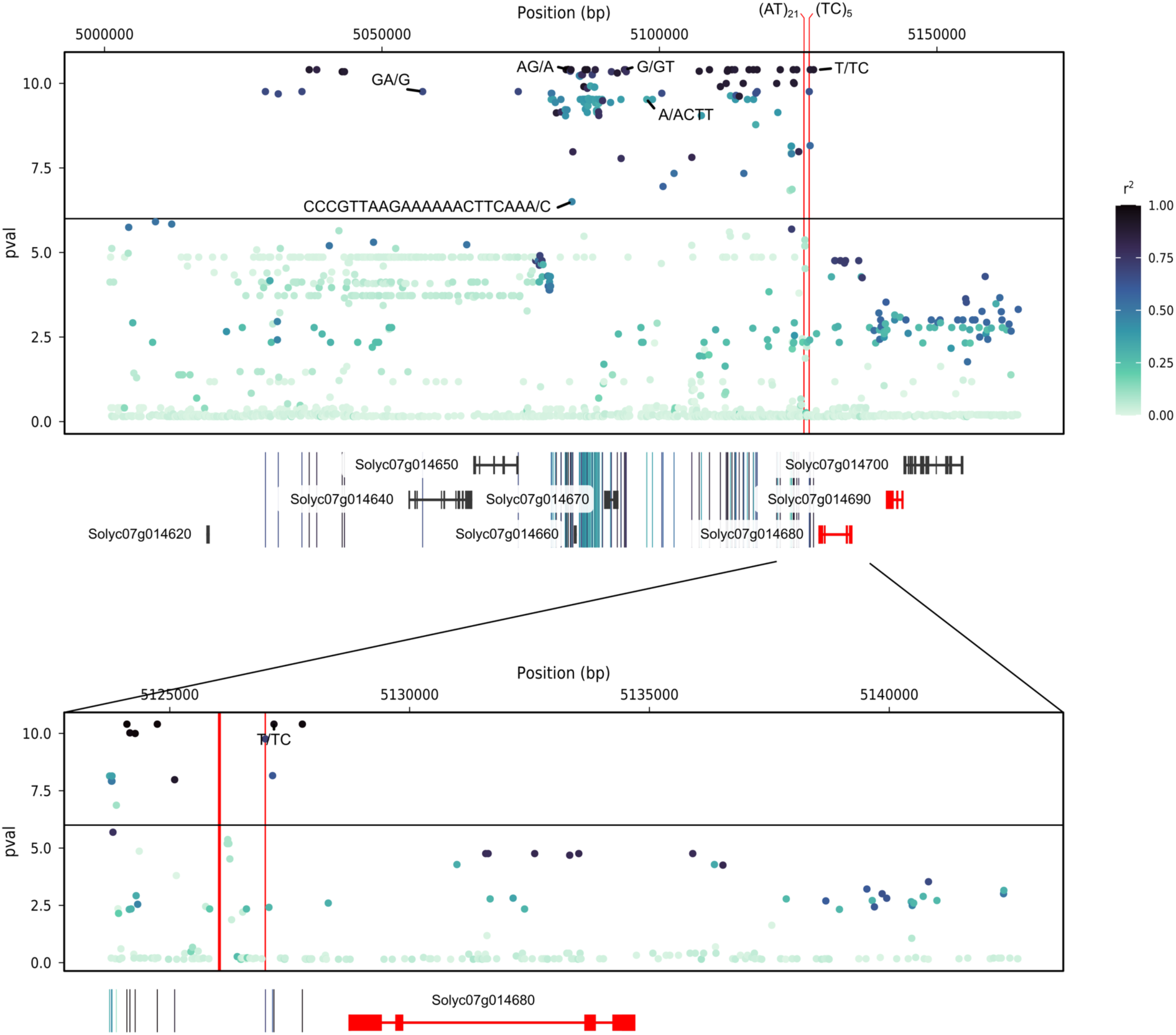
**Local visualisation of GWAS analysis for *Sl*HKT1.2 root expression** The upper panel shows the local Manhattan plot of the GWAS results. The colour legends correspond to the linkage disequilibrium (r^2^) between the most significant markers and all other positions. Short indels are labelled only if they are significant for GWAS. The red genes highlight *Sl*HKT1.2 (Solyc07g014680) and *Sl*HKT1.1 (Solyc07g014690). The position of two short tandem repeats (not analysed by GWAS) is represented by the red vertical lines. The lower panel show the annotated genes (SL4.0 genome version) in the region below the peak.

**Table 2.** Promoter sequence polymorphisms of *Sl*HKT1.2.

| Genome position | Position from TSS | Allele (ref/alt) | GWAS *p*-value | Cis-regulatory element | Biological function |
| --- | --- | --- | --- | --- | --- |
| 5,127,764 | -972 | C/T | **5,10E-11** | Ref : (+)<br>**ASF1MOTIFCAMV**<br>-975 TGA**C**G -971<br>Alt allele = No motif | - “**ASF-1 binding site**” in CaMV 35S promoter found in many promoters<br>- Are involved in transcriptional activation of several genes by auxin and/or salicylic acid |
|  |  |  |  | Ref : (+) WRKY71OS<br>-975 TGA**C** -970<br>Alt allele = No motif | - “**A core of TGAC-containing W-box**” of, e.g., Amy32b promoter<br>- Binding site of rice WRKY71, a transcriptional repressor of the gibberellin signalling pathway |
| 5,127,172 | -1,564 | T/TC | **5,10E-11** | REF : (-)<br>DPBFCOREDCDC3<br>-1565 T(-)CGTGTG -1558<br>Alt allele = No motif | - ABA- responsive element<br>- ABRE/CAMTA motif<br>- Present in the upstream region of *OsSOS1* |
| 5,127,143 | -1,593 | A/C | **1,37E-08** | No differentially affected motif |  |
| 5,126,994 | -1,742 | G/T | **6,02E-10** | No differentially affected motif |  |

This table displays all significant polymorphisms in the *SIHKT1.2* promoter sequence (-2000: 0 bp of TSS). Cis-regulatory elements were identified using the PLACE database (https://www.dna.affrc.go.jp/PLACE/?action=newplace). ‘Genome position’ corresponds to the true physical position according to SL4.0. ‘Position from TSS’ corresponds to the position of the mutation according to zero, the position of the Transcription Start Site.

To further contextualise these findings, we investigated whether the candidate polymorphisms were located in evolutionarily conserved regions of the promoter. For this purpose, we aligned the promoter sequences of *Sl*HKT1.2 homologs of ten wild tomato cross-compatible species – *S. galapagense*, *S. pimpinellifolium*, *S. chmielewskii*, *S. neorickii*, *S. corneliomulleri*, *S. peruvianum*, *S. chilense*, *S. pennellii*, *S. habrochaites* and *S. lycopersicoides*. We then assessed the level of conservation by measuring the allelic frequency of the alternative alleles (**Supplementary Figure 2**). Our analysis revealed that the *Sl*HKT1.2 promoter sequence was highly conserved up to position −1,800 relative to the TSS in these species. Furthermore, among the four candidate polymorphisms, two (−1,593 and −1,564) were conserved across all species. In particular, the ABRE motif, whose disruptive variant was suspected of being responsible for the differential expression of the *Sl*HKT1.2 loci, was perfectly conserved in all the species examined.

### 3.5. Diversity and evolutionary history of HKT1.2

To gain insights into the evolutionary history of HKT1.2 alleles identified in our study, we performed a large-scale genomic analysis. In order to construct a global perspective of the distribution and demographic histories of the HKT1.2 alleles, we integrated into our core collection data an additional set of 163 genome sequences sourced from the Varitome project [24]. This expanded dataset includes genomes from *S. pimpinellifolium* (SP*), S. lycopersicum var. cerasiforme* (SLC), and *S. lycopersicum var. lycopersicum* (SLL), thereby capturing the genetic diversity at the epicentre of tomato origin of domestication. Interestingly, the allele associated with a high HKT1.2 expression is absent from the *S. pimpinellifolium* background. Its increasing prevalence in domesticated forms of tomato (piecharts are arranged in order according to the tomato domestication model proposed by [41]) suggested that the highly expressed HKT1.2 allele underwent selection during the domestication process (**Supplementary figure 3.a**). To corroborate these findings, we also examined the Extended Haplotype Homozygosity (EHH) decay patterns surrounding the HKT1.2 gene. Our analysis suggested that the ‘cerasiforme’ haplotype underwent artificial selection during the course of tomato domestication. Specifically, the derived haplotype (more frequent in domesticated forms) exhibited a slow decay in EHH values, corresponding to a recent selection event affecting the polymorphisms in proximity to the HKT1.2 derived allele (**Supplementary Figure 3.b**).

### 3.6. Detection of Na^+^ content QTL under hybrid background

To assess the potential of the QTLs detected on the CC panel containing only inbred lines for breeding applications, the greenhouse trials were replicated using an hybrid population (Test-cross: TC) obtained by crossing each line of the core collection with a large-fruited tomato [14]. As expected from a test-cross panel, all the traits measured showed a significantly reduced coefficient of variation (**Supplementary Table 3**). Analysis of correlations between mineral traits measured on the CC and TC panels showed that all Na^+^ and K ^+^content traits were correlated with each other, ranging from *r*=0.46 for fruit K^+^ to *r*=0.61 for Na^+^ petiole content (**Supplementary Figure 4**). For Na^+^ leaf petiole content, fruit Na^+^ content and K^+^ petiole content, the major QTL on chromosome 7 was also detected (**Figure 5**). The lead SNP was not identical to that identified in the core collection, probably because the composition of the two panels was slightly different (**Supplementary Table 3**), causing an imbalance in allele frequencies. Also, the percentage of variance explained was significantly lower than for QTL identified using the core collection (PVE = 0.11 for petiole Na^+^ content in the TC panel). These results suggested that HKT1.2 QTL also had an effect at the heterozygous state, making it interesting for F1 tomato breeding. We also detected QTLs specific to the test-cross panel which had putative candidate genes. For instance, for petiole K^+^ on chromosome 1, the lead marker in position 77,939,937 was near a K^+^ efflux antiporter (Solyc01g094290) mostly expressed in leaf tissues.

**Figure 5.**
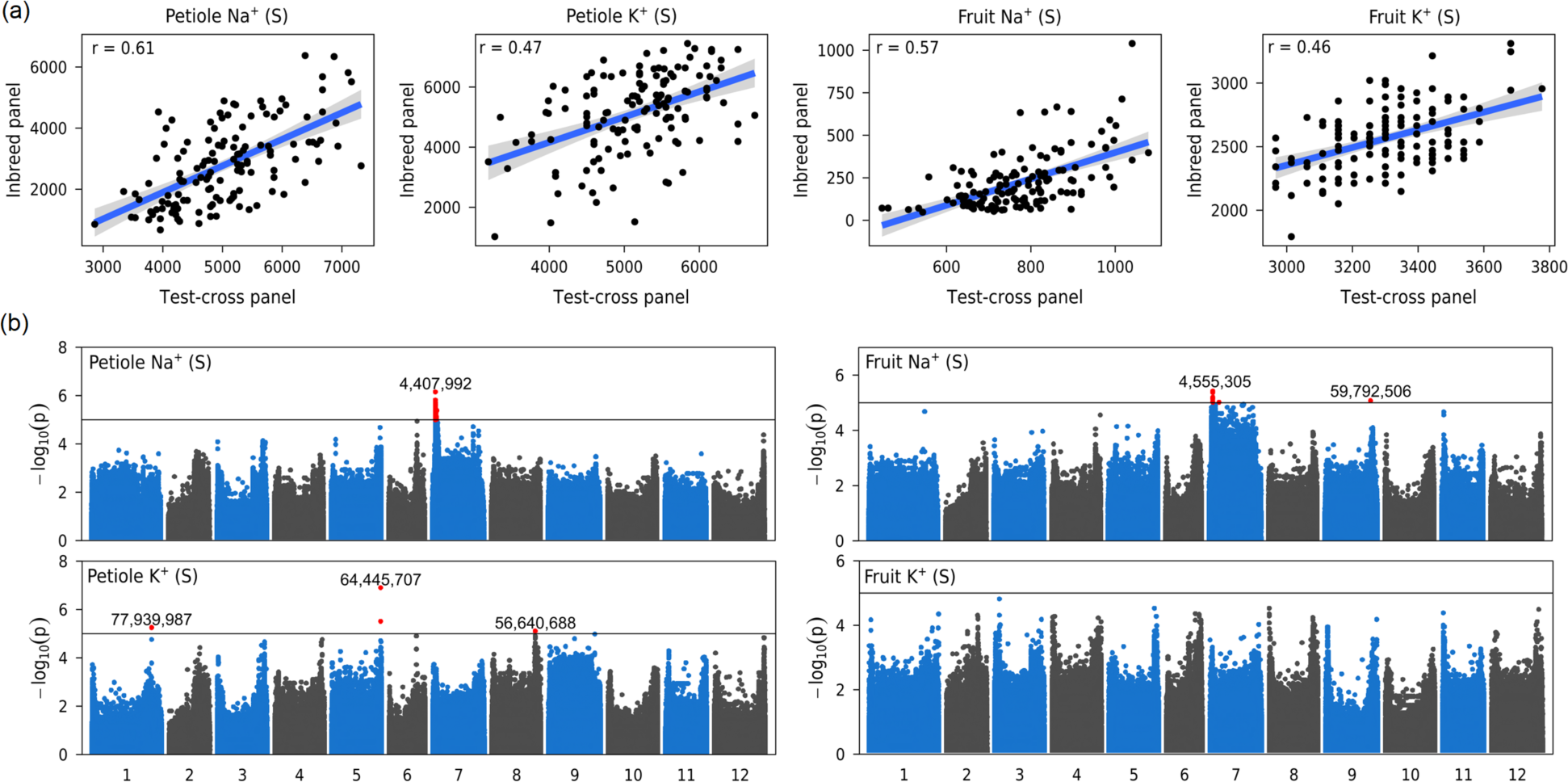
**GWAS for Na^+^ and K^+^ content in an hybrid genetic background** (**a**) Relationships between traits measured in the core collection (Inbred panel) and the test-cross panel and Pearson correlation coefficient (**b**) Manhattan plot showing marker-trait associations for Petiole Na^+^ and K^+^ in stress condition (S) and Fruit Na^+^ and K^+^ in stress condition (S) in the Test cross panel.

## 4. Discussion

### 4.1. Is sodium content a reliable marker of tomato salt tolerance?

Some wild relatives of tomato have higher salt tolerance than cultivated tomato and can therefore provide alleles for genetic improvement of salinity tolerance [6,11]. This requires the identification of interesting germplasm and traits implicated in stress tolerance before the identification of QTLs. Salt tolerance is usually assessed as the percent biomass production in saline versus control conditions. However, evaluating salinity tolerance strictly on the basis of biomass differential offers limited understanding of the underlying physiological mechanisms. A more fruitful approach lies in investigating the genetic control of specific downstream traits that contribute collectively to the overall salt tolerance. Among them, the one that is undoubtedly most described is sodium exclusion. In fact, in several species, a negative correlation between leaf sodium content and salinity tolerance have been described [42,43]. In tomato, whether or not this relationship exists is unclear : several studies didn’t find any correlation between foliar sodium content and salinity tolerance [44,45]. In our study, we didn’t find any significant correlation between biomass-based salt tolerance and shoot sodium content at the juvenile stage. However, at the adult stage, we observed a negative correlation between sodium content and biomass proxies (stem diameter, leaf length and plant height). Thus, the relationship between leaf Na^+^ content and salt tolerance might be detected only at mature stage. This effect has been documented in monocotyledons: some authors showed that maize and wheat cultivars with different salt tolerance can grow at the same rate for two to four weeks, before revealing a difference in growth kinetics [4]. The kinetics of ion build-up in shoots to toxic concentrations is, however, not well documented in solanaceous species (and is also probably genotype-dependent). However, it is highly likely that the situation is similar in the case of tomato. If this is the case, screening plants for salt tolerance at too early a stage is akin to phenotyping for osmotic stress tolerance. However, as we have seen, genotypic differences in sodium accumulation are detectable at the early stage, even under control conditions (without exogenous addition of sodium to the fertigation solution).

Still, the relevance of leaf sodium content as a tolerance trait *per se* is open to debate. For instance, salt-tolerant wild relatives of tomato, such as *S. chilense* and *S. pennellii*, exhibit significantly higher leaf Na^+^ content compared to their more sensitive counterpart, *S. lycopersicum* - up to ten-fold higher in the case of *S. chilense* [5,7]. These species seem to have a different strategy from *S. lycopersicum*, maintaining high levels of sodium leaf content, but investing more in sodium sequestration as suggested by an increased expression of tonoplast-localised NHX-type Na^+^/H^+^ exchangers compared to their homologues in cultivated tomato [5,7]. In such species, the high accumulation of Na^+^ in the leaves acts as a cheap osmoticum to minimise the high energy cost of osmolyte synthesis. This raises the question of whether leaf Na^+^ content should be considered as a target trait for breeding salt-tolerant tomato varieties? Our data suggest a positive answer, at least in the genetic background of *S. lycopersicum*; this is further supported by studies showing that knock-down of genes responsible for sodium ion translocation from roots to leaves, in particular *Sl*HKT1.2 and *Sl*SOS1, results in increased sensitivity to salt stress [12,46]. However, the introgression of exotic genetic material from ‘includer’ species (*S. chilense*, *S. pennellii*) could compromise this assertion, a point that remains to be verified.

### 4.2. Genetic diversity of above-ground sodium content is controlled by a few QTLs

During GWAS analysis, we identified a limited number of QTLs for Na^+^ and K^+^ content levels. Notably, excluding the hotspot QTL located on chromosome 7, only a 13 QTL were detected. This finding is consistent with existing literature on the subject. For example, a recent GWAS study conducted on a panel of 250 *S. pimpinellifolium* accessions for Na^+^ content in leaves also reported a little number of QTLs. Specifically, only three QTLs were detected and, one of these QTLs corresponded closely to the QTL identified in our own study on chromosome 4 [47]. Similarly, in the Wang study [11], no significant association between markers and sodium or potassium levels at shoot level was found. The QTL found on chromosome 7 is not a novel finding. It was reported by Villalta et al., [10] who analysed leaf and stem sodium content in two different RIL populations derived from the cross of a cultivated tomato accession (SLL) with two wild relatives (*S. pimpinellifolium* and *S. galapagense*). The identification of the underlying candidate genes (HKT1.1 and HKT1.2) was published a few years later by the same team [15]. Analysis of the knock-down of each HKT1 gene was conducted using two near-isogenic lines (NILs) that were homozygous for either the *S. lycopersicum* allele or the *S. galapagense* allele and the authors showed that the knock-down of HKT1.2 gene from *S. lycopersicum* and S. *galapagense* increased the leaf Na^+^/K^+^ ratio [48] and had a significant negative impact on yield [49]. The effect of HKT1.1, however, is less clear. Indeed Jaime-Pérez, N. et al. (2017) [48] showed that silencing of *Sl*HKT1.1/*Sc*HKT1.1 resulted in a non-significant effect while Romero-Aranda et al., [49] observed that the knockdown of *Sc*HKT1.1 resulted in the reduction of the Na^+^/K^+^ ratio (but still no significant effect for *Sl*HKT1.1). However, these studies have not been able to determine which gene is at the source of the chromosome 7 QTL.

### 4.3. The sodium content QTL on chromosome 7 is characterised by a variation in *Sl*HKT1.2 gene expression

The search for polymorphisms in the promoter region of *Sl*HKT1.2 did not reveal any obvious candidate polymorphisms (e.g. large deletions in the promoter) which could explain the variation in gene expression. However, several variants significantly associated with a difference in HKT1.2 expression have been detected in the promoter sequence: for example, an insertion of one base (T/TC) at 900 bp from the initiation start) was significantly associated with the change in gene expression and could be a good candidate modification for functional validation through prime editing. In addition, we found a large deletion in the *de novo* assembly sequence of *Sl*HKT1.2 corresponding to a tandem repeat (3000 bp). As tandem repeats are known to affect the transcription of neighbouring genes by regulating chromatin structure and composition, this structural variant could be an other interesting candidate modification. Also, in *Arabidopsis*, it was established that a deletion in a tandem repeat sequence approximately 5 kb upstream of At*HKT1* is responsible for the reduced root expression of At*HKT1* [50]. Unfortunately, we were not able to verify if this deletion was present in accessions with only low expression as read-alignment failed for most of the accessions due to short read sequencing. From an evolutionary perspective, the absence of this allele in the *S*. *pimpinellifolium* background and its increasing prevalence in domesticated forms of tomato suggest that the highly-expressed HKT1.2 allele was indeed selected during domestication. However, the evolutionary mechanism behind this process is still unclear, especially as other salinity tolerance QTLs such as *Sl*HAK12 and *Sl*SOS1 were lost during domestication [12]. Sodium, at low levels, is beneficial to tomato growth and yield [51], especially under suboptimal potassium supply; we may thus hypothesise that the fixation of an highly expressed *Sl*HKT1;2 allele could have occurred during the domestication of tomato and was then indirectly selected for growth under optimal conditions.

## 5. Conclusion

The results of the genome-wide association study offer a comprehensive list of promising QTL and candidate alleles that deserve further exploration to understand the genetic and molecular mechanisms underlying variability in mineral content under salt stress conditions. In particular, we have identified a major QTL located on chromosome 7 colocalising with an eQTL for HKT1.2 gene expression in root, making this gene a prime candidate for further research. A natural variation identified in the *Sl*HKT1.2 promotor, which has undergone selection during tomato domestication and improvement, holds great promise for enhancing tomato salt tolerance and represents an important target for the development of salt-tolerant tomato germplasm.

## Data availability

Raw genome reads corresponding to the accession sequences from the core-collection can be found in NCBI (https://www.ncbi.nlm.nih.gov/sra) database under accession number PRJNA1014227. All other data supporting the findings of this study are available in the supplementary data published online.

## Supplementary Material

**Supplementary Table 1.** List of the accessions of the diversity panel.

**Supplementary Table 2:** Mean phenotypic values measured under control (C), stress (S) or plasticity (D) condition for the inbred core-collection at the young stage and in stress condition at the adult stage

**Supplementary Table 3:** Mean phenotypic values measured under stress (S) condition for the test-cross core-collection

**Supplementary Table 4**: Mean root expression of SlHKT1.1 and SlHKT1.2 (expressed in fold change)

**Supplementary Table 5**: Summary of QTLs detected for salt tolerance-related traits

**Supplementary Figure 1**. Expression profile of candidate genes identified through GWAS in various tissues

**Supplementary Figure 2.** Conservation profile of the HKT1.2 promoter sequence in 9 wild tomato relative species

**Supplementary Figure 3.** QTL frequency in different tomato genetic groups and Haplotype Homozygosity Decay

**Supplementary Figure 4.** Hierarchical clustering heatmap of Pearson correlation coefficients between Na/K content, fruit weight and yield in the test-cross panel

## Acknowledgement

We are thankful to Mélanie Andrin, Anaïs Roger, Matilde Van-Ommen and Camille Rabeau for taking care of the plants and theirs help for plant phenotyping, to the vegetable resources centre (CRBLeg) of GAFL for keeping the seeds of the CC panel available, to the greenhouse staff of UE A2M “Arboriculture et Maraichage Méditerranéens” and to Sandrine Chay (INRAE, UMR BPMP / SAME plateform) for ionomic analysis. This work was supported by the ERA-NET SusCrop ROOT (Grant No. 771134) and PRIMA-VEG-ADAPT (Grant No. 01DH19019) projects.

## Author contribution

MC and AH: conceptualization, data analyses and writing the manuscript; YC, EP, RD, MG, CG: data acquisition; FB bioinformatic data analysis; MC, RK, CT and RS : funding acquisition.

