## Supplementary figures for "Characterization of a major QTL for sodium accumulation in tomato shoot"

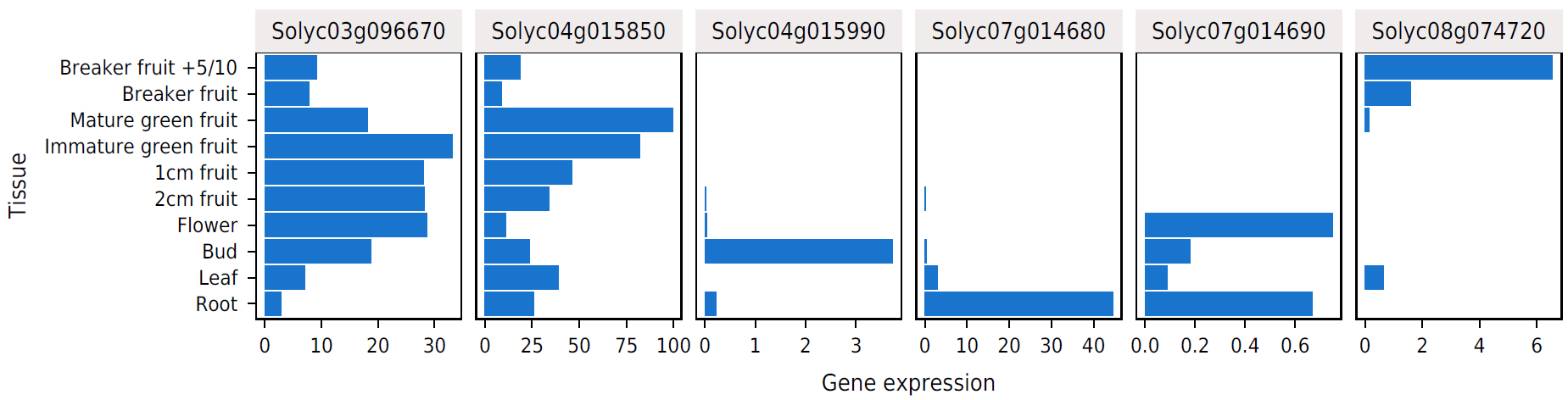

**Supplementary Figure 1**. **Expression profile of candidate genes identified through GWAS in various tissues**

Public data from Solanum lycopersicum cv. ‘Heinz 1706’ (The Tomato Genome Consortium 2012)

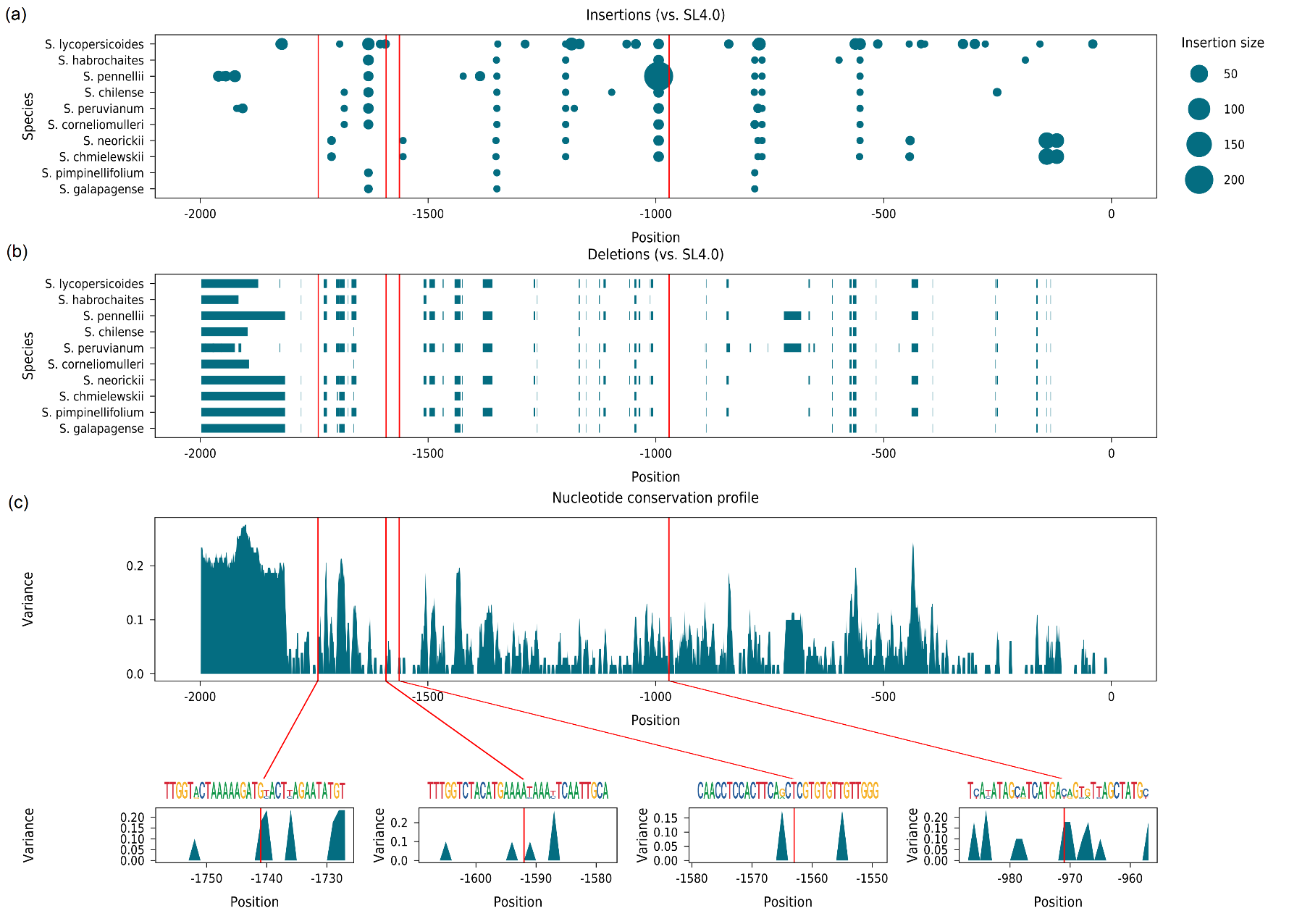
**Supplementary Figure 2.** Conservation profile of the HKT1.2 promoter sequence in 9 wild tomato relative species

Each plot shows the alignment of the wild relative species to the SL4.0 reference genome. Dot size is proportional to the size of the length of insertions. Red lines indicate the position of the four significant SNPs for GWAS analysis of SlHKT1.2 root expression. (b) Position of deletions in wild relatives (‘real size’). (c) Nucleotide conservation profile measured as the variance of allele numbers for a given position. (d) Zoom on candidate polymorphisms and sequence logo displaying relative base frequency.

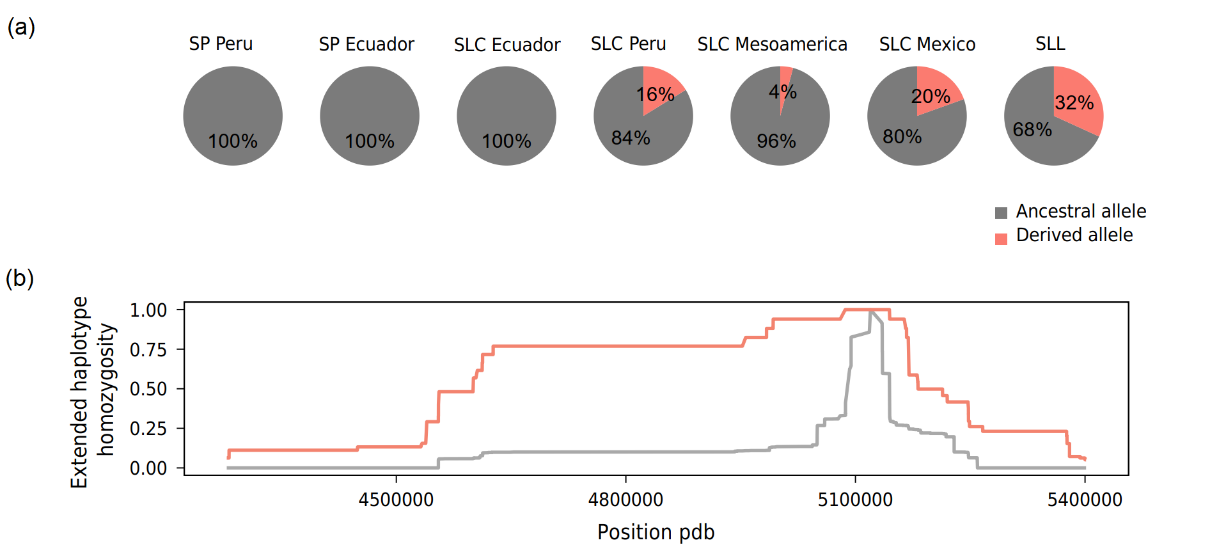

**Supplementary Figure 3.**  **QTL frequency in different tomato genetic groups and Haplotype Homozygosity Decay**

(**a)** QTL frequency in different genetic groups of the extended panel (SP *S. pimpinellifolium*: - SLC: *S. lycopersicum var. cerasiforme*; SLL: *S. lycopersicum var. lycopersicum*). Numbers represent the percentages of ancestral (grey) and derived alleles. (**b**) Extended Haplotype Homozygosity (EHH) decay of *Sl*HKT1.2. Grey and red lines represent EHH decay of ancestral (SL4.00 reference allele) and derived alleles, respectively.

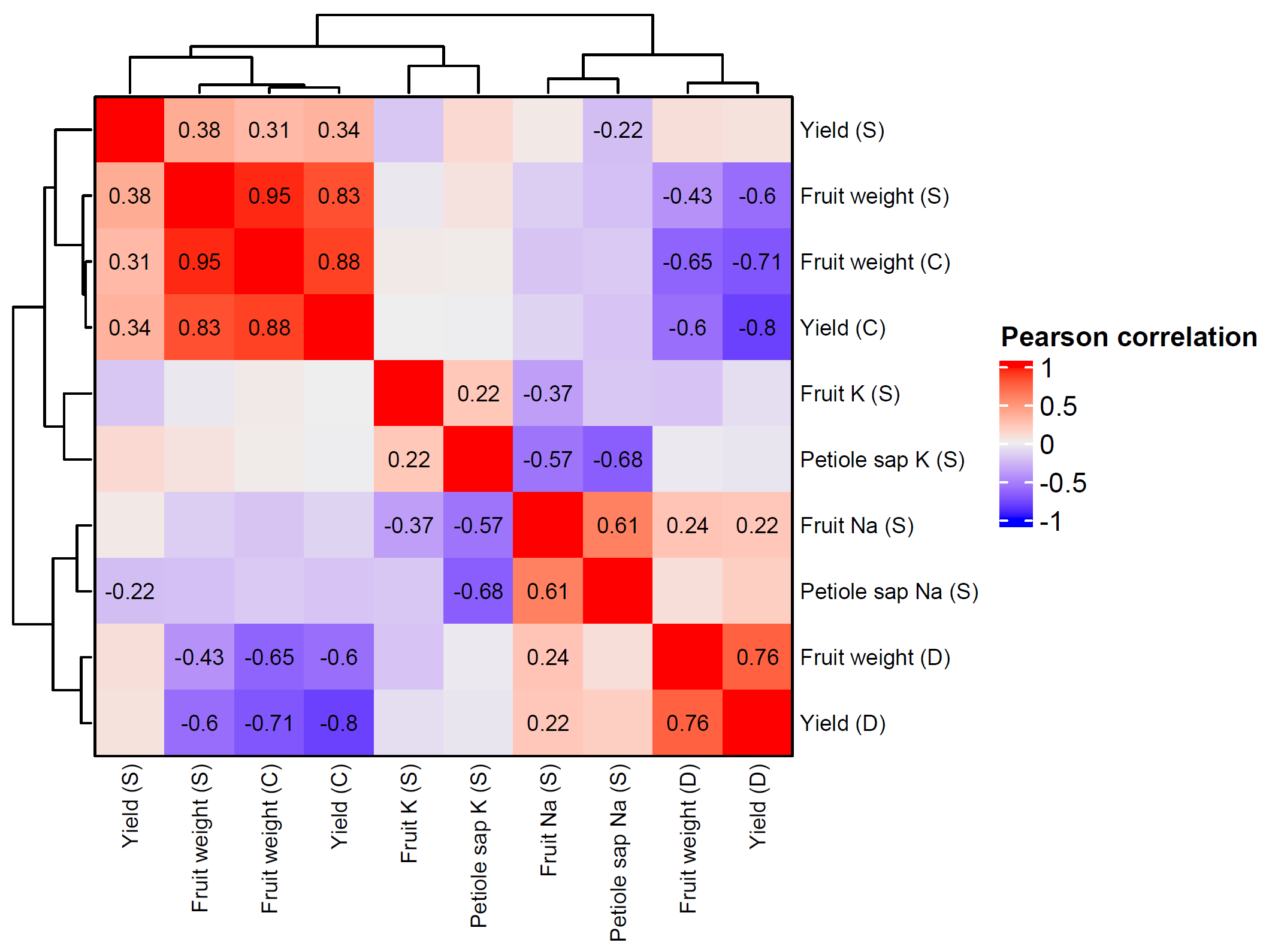

**Supplementary Figure 4. Hierarchical clustering heatmap of Pearson correlation coefficients between Na/K content, fruit weight and yield in the test-cross panel**

This matrix represents the recorded traits under three conditions: Control (C), Salinity Treatment (S), and the ratio of the two treatments (D). The colour gradient provides insight into the correlation type; a value of 1 signifies a complete positive correlation (represented by dark red), while a value of -1 indicates a full negative correlation (represented by dark blue) between two traits. Only those correlations that are statistically significant (p-value < 0.01) have been shown.
